# Mitochondrial protein FgDML1 impacts DON toxin biosynthesis and cyazofamid sensitivity in *Fusarium graminearum* by affecting mitochondrial homeostasis

**DOI:** 10.1101/2025.05.23.655648

**Authors:** Chenguang Wang, Xuewei Mao, Weiwei Cong, Lin Yang, Yiping Hou

## Abstract

*Fusarium graminearum* is a global pathogen responsible for Fusarium head blight (FHB) in wheat, causing substantial yield losses and producing the mycotoxin deoxynivalenol (DON), which poses a threat to both human and animal health. *Drosophila melanogaster* Misato-Like protein (DML1) plays a critical role in regulating mitochondrial function, yet its function in filamentous fungi remains unexplored. In this study, we characterized FgDML1 in *F. graminearum*. FgDML1 interacts with the mitochondrial fission and fusion protein FgDnm1 to maintain mitochondrial stability, thereby positively regulating acetyl-CoA levels and ATP synthesis, which influences toxisome formation and ultimately affects DON toxin biosynthesis. Additionally, FgDML1 is involved in the regulation of toxin biosynthetic enzyme expression. In the ΔFgDML1 mutant, Complex III enzyme activity decreased, overexpression of complex III assembly factors FgQCR2, FgQCR8, and FgQCR9 may induce conformational changes in the Qi-site protein, specifically altering the sensitivity of *F. graminearum* to respiratory inhibitor cyazofamid not Qo-site inhibitor pyraclostrobin and other fungicides. Furthermore, the loss of FgDML1 leads to defects in nutrient utilization, as well as in asexual and sexual reproduction, and pathogenicity. In conclusion, this study identifies a novel regulatory role for FgDML1 in DON toxin biosynthesis and cyazofamid sensitivity in *F. graminearum*. Our study provides a theoretical framework for understanding DON biosynthesis regulation in *F. graminearum* and identifies potential molecular targets for FHB control.

## Introduction

Fusarium head blight (FHB), caused by *Fusarium graminearum*, is a devastating fungal disease that affects cereal crops, leading to significant yield losses in wheat-growing regions worldwide(Dean et al., 2012). This disease is particularly concerning due to the production of various mycotoxins, including trichothecenes and zearalenone, during fungal infection(Hai et al., 2023). Among these, deoxynivalenol (DON), a type of trichothecene, is the most commonly detected mycotoxin(Merhej et al., 2011). The DON toxin poses a serious threat to both human and animal health, significantly compromising immune and reproductive systems(Knutsen et al., 2017). Chronic exposure to DON can lead to anorexia, neurological disorders, and immune dysregulation, while acute high-dose exposure may result in vomiting, diarrhea, neurological disturbances, and even miscarriage or stillbirth(Sumarah, 2022; Zhang et al., 2024). Studies have shown that increased DON levels are positively correlated with the pathogenicity rate of *F. graminearum*(Bönnighausen et al., 2019; Ilgen et al., 2009; Mudge et al., 2006). As FHB continues to spread and its severity increases, DON contamination is emerging as a critical global concern(Duan et al., 2020; Liu et al., 2016). This highlights the need for effective strategies to manage and mitigate the impact of both the disease and the associated mycotoxin contamination.

The absence of resistant cultivars necessitates reliance on chemical methods as the primary approach for controlling FHB(Liu et al., 2019). However, the inappropriate application of fungicides has led to the emergence of resistance, particularly against carbendazim, tebuconazole, and pyraclostrobin(Andrade et al., 2022; Yi et al., 2023; Zhou et al., 2024). Notably, carbendazim has been demonstrated to stimulate the biosynthesis of DON toxin, especially in fungicide-resistant populations of *F. graminearum*(Li et al., 2019; Zhou et al., 2020). Similarly, pyraclostrobin significantly promotes DON biosynthesis, with no selectivity between resistant and sensitive strains. Although tebuconazole has not been directly implicated in inducing DON production, it is classified as a fungicide with a moderate risk of resistance development(Duan et al., 2020). Field observations have revealed diverse resistance mechanisms to tebuconazole, complicating management efforts(Liu et al., 2011; Zhao et al., 2021). The potential evolution of resistance to tebuconazole is particularly concerning, as it could undermine its efficacy and exacerbate the challenge of controlling DON contamination. There is an urgent need for a fungicide that has not yet been used in *F. graminearum* to address the DON toxin contamination caused by fungicide resistance. Cyazofamid is a mitochondrial respiratory chain complex III inhibitor with demonstrated efficacy in controlling fungal diseases. It specifically binds to the Qi site of complex III, blocking the reduction of ubiquinone and inhibiting ATP synthesis, thereby exerting its fungicidal effect(Li et al., 2014; Mitani et al., 2001). However, it remains unclear whether other proteins within the intricate regulatory network of the respiratory chain play a role in modulating fungal sensitivity to cyazofamid. Further investigations are needed to elucidate the potential involvement of these regulatory components.

The synthesis of DON toxin is primarily regulated by genes involved in trichothecene biosynthesis, known as *FgTRIs*. Many of these *FgTRI* genes have been well characterized(Merhej et al., 2011). The initial step in DON synthesis is catalyzed by the trichothecene synthase *FgTri5*, which facilitates the cyclization of farnesyl pyrophosphate (FPP) to form trichodiene (TDN)(Kimura et al., 2007). However, before this catalytic process can occur, the organism must first obtain sufficient ATP from the mitochondria. ATP is critical not only for the basic cellular functions but also for driving several key reactions in the isoprenoid biosynthesis pathway. Specifically, ATP is required for the enzymatic conversion of precursors, leading to the production of FPP, a vital intermediate in the trichothecene biosynthesis pathway. The availability of ATP therefore directly impacts the efficient synthesis of FPP, which, in turn, regulates the downstream production of DON toxin. Without an adequate supply of ATP, these biosynthetic pathways cannot proceed efficiently, thus hindering the synthesis of both the precursor molecules and the final toxin product(Alexander et al., 2009). Pathogen-intrinsic ATP homeostasis is recognized as a critical, rate-limiting determinant for toxin biosynthesis. Previous studies indicate that dual-target inhibition of ATP synthase (AtpA) and adenine deaminase (Ade) by a specific small-molecule probe effectively depletes intracellular ATP, consequently suppressing the synthesis of key virulence factors TcdA and TcdB transcriptionally and translationally(Marreddy et al., 2024). The systemic toxicity of Anthrax Edema Toxin (ET) is primarily attributed to its catalytic activity, which depletes the host cell’s ATP reservoir, thereby triggering a bioenergetic collapse that culminates in cell lysis and death(Liu et al., 2025). The biosynthesis of DON entails a reorganization of the endoplasmic reticulum into a specialized compartment termed the "toxisome"(Tang et al., 2018). The assembly of the toxisome coincides with the aggregation of key biosynthetic enzymes, which in turn enhances the efficiency of DON production. Concurrently, this compartmentalization serves as a self-defense mechanism, protecting the fungus from the autotoxicity of TRI pathway intermediates (Boenisch et al., 2017). The proteins TRI1, TRI4, TRI14, and Hmr1 are confirmed constituents of this structure(Kistler and Broz, 2015; Menke et al., 2013).

### Drosophila melanogaster

Misato-Like protein (DML1) encodes a Misato-like protein in *Saccharomyces cerevisiae*, an essential factor involved in mitochondrial DNA (mtDNA) inheritance. In addition to its role in mtDNA maintenance, DML is implicated in mitochondrial compartmentalization and chromosome segregation. The protein shares structural features with members of the GTPase family, suggesting potential involvement in GTP-dependent processes(Gurvitz et al., 2002). In *Homo sapiens*, the DML protein is localized to the mitochondria, where silencing of DML results in a transition from a networked mitochondrial morphology to a fragmented form. This indicates that DML plays a critical role in mitochondrial distribution, morphological fusion, and overall mitochondrial dynamics(Kimura and Okano, 2007). In prokaryotes, the homologous protein FtsZ, which shares functional similarities with DML, has been identified as a target for antimicrobial agents(Huang et al., 2007; Vollmer, 2006). However, despite these insights from studies in the aforementioned organisms, the exact role of DML in filamentous fungi remains undetermined.

This study systematically characterized the functional roles of DML protein in *F. graminearum*(FgDML1), including mycelial growth, asexual development, and sexual reproduction. More importantly, we found that FgDML1 reduces DON toxin synthesis by decreasing the enzyme activity of complex III, reducing ATP synthesis, and lowering acetyl-CoA accumulation. Furthermore, the deletion of FgDML1 results in compensatory upregulation of complex III assembly factors, which may alter the conformation of the Qi site protein and subsequently modulates sensitivity to cyazofamid.

## Results

### Identification, deletion, and complementation of FgDML1

The DML1 protein from S. cerevisiae was used as a query for a BLAST search against the Fusarium genome database, resulting in the identification of the putative DML1 gene FgDML1 (FGSG_05390) in *F. graminearum*. The FgDML1 gene is 1455 bp in length, contains three introns, and encodes a 484-amino acid protein with 23.88% homology to its counterpart in S. cerevisiae. Phylogenetic analysis was performed using DML1 sequences from other species obtained via NCBI, and a phylogenetic tree was constructed (Fig. 1A). Structural analysis revealed the presence of the Misat_Tub_SegII and Tubulin_3 domains. Furthermore, subcellular localization studies confirmed that FgDML1 localizes to mitochondria, as demonstrated by colocalization with a mitochondria-specific dye MitoTracker Red CMXRos (Fig. 1B).

**FIG. 1.**
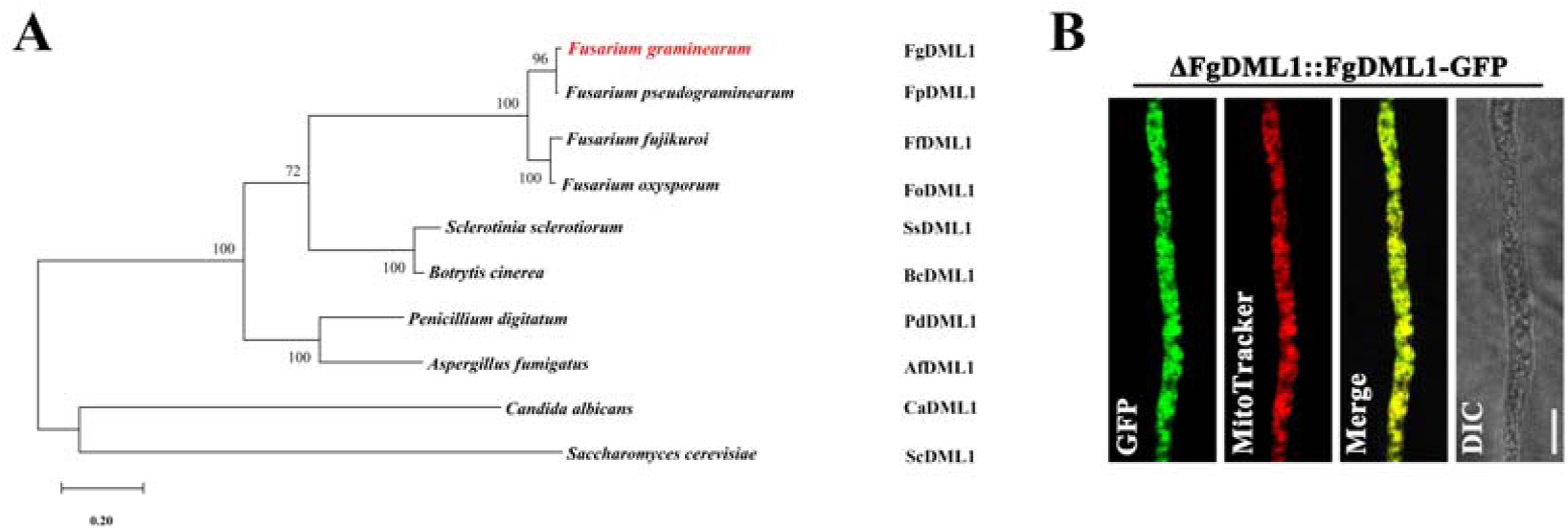
Identification and localization of FgDML1 in *Fusarium graminearum*. (A) The phylogenetic tree was constructed based on the amino acid sequences of DML1 homologs from 10 species using the maximum likelihood method in Mega X. Bootstrap values from 1000 replications were displayed on the branches. (B) FgDML1 was localized to the mitochondria.

To investigate the function of FgDML1, ΔFgDML1 deletion mutant was constructed and validated via PCR and Southern blot analysis (Fig. S2), confirming the mutant as a single-copy knockout. To ensure that the observed phenotypes were specifically attributable to the deletion of FgDML1, an in situ complemented mutant, ΔFgDML1-C, was generated to restore FgDML1.

### FgDML1 regulates vegetative growth

To assess the role of FgDML1 in vegetative growth, PH-1, ΔFgDML1, and ΔFgDML1-C were cultured on various media, including PDA, CM, MM, and V8, to measure growth rates. The ΔFgDML1 strain displayed a distinct growth phenotype characterized by retardation in radial growth and the formation of more compact, denser hyphal networks on all tested media compared to the PH-1 and ΔFgDML-C strains. (Fig. 2A). Moreover, ΔFgDML1 displayed pronounced hyphal curvature, which was not observed in PH-1 or ΔFgDML1-C (Fig. 2B). These findings demonstrate that FgDML1 is a positive regulator of virulence in *F. graminearum*.

**FIG. 2.**
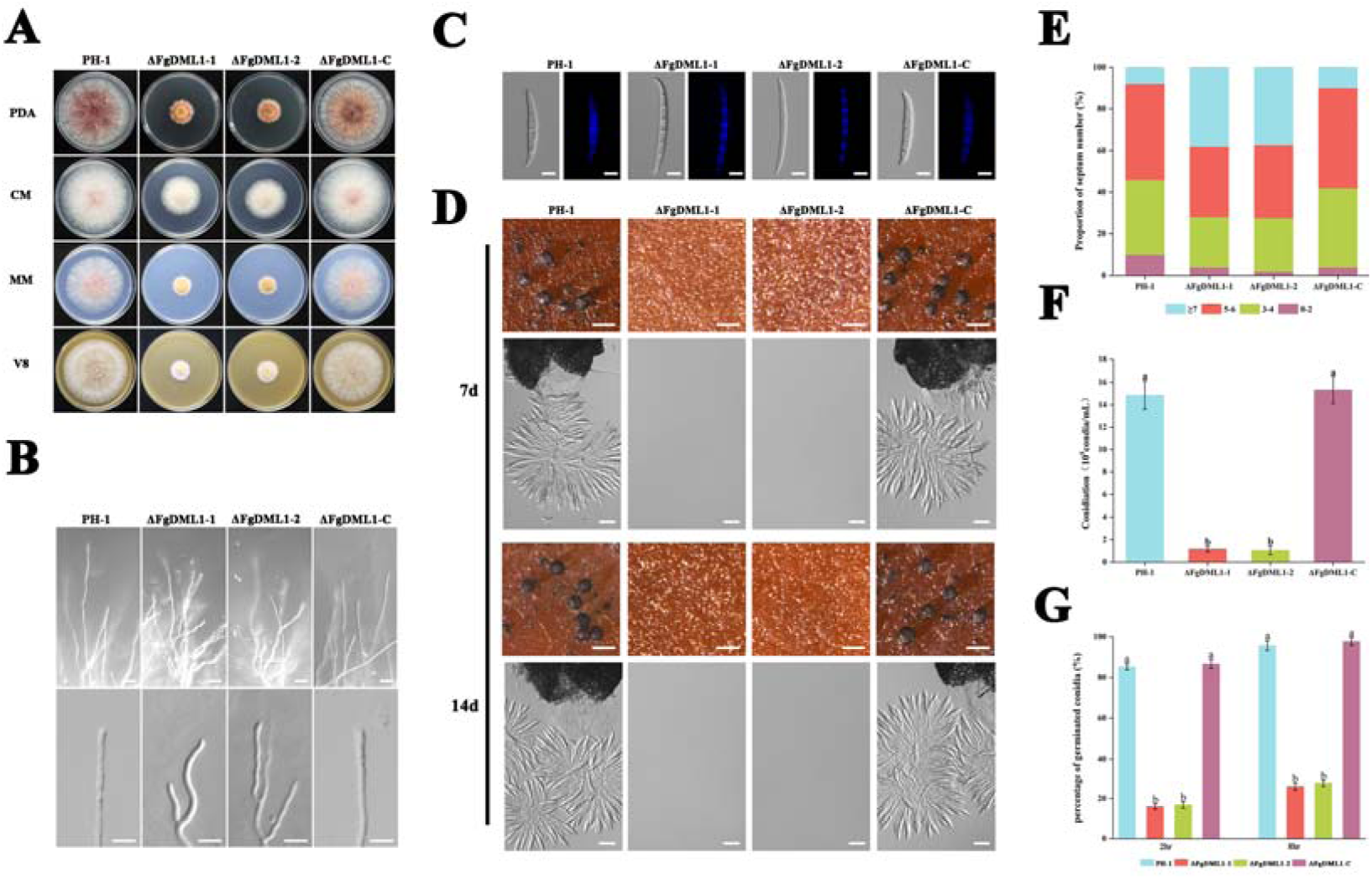
Effects of ΔFgDML1 on asexual and sexual development. (A) Mycelial growth rate. PH-1, ΔFgDML1, ΔFgDML1-C were cultured for 3 days on PDA, CM, MM, and V8 solid media at 25°C. (B) Mycelial morphology. The mycelial morphology of PH-1, ΔFgDML1, and ΔFgDML1-C was examined on WA plates. Scale bar = 72 μm. (C) Conidial length. Conidia of ΔFgDML1 were significantly longer in length compared to PH-1 and ΔFgDML1-C. Conidia were germinated in MBL medium, stained with Calcofluor White (CFW), and observed using an Olympus IX-71 microscope. (D) Sexual reproduction. PH-1 and ΔFgDML1-C produced perithecia and asci containing ascospores after 7 or 14 days of incubation on CA medium, while the ΔFgDML1 failed to produce these structures under the same conditions. Perithecia and ascospores were observed using a Nikon SMZ25 fluorescent stereo microscope and an Olympus IX-71 inverted fluorescence microscope. Top image: perithecium, scale bar = 1 mm; bottom image: ascospores, scale bar = 50 μm. (E) Proportion of conidial septa. The proportion of conidial septa was assessed in the tested strains. (F) Conidial production. Conidia were harvested after 4 days of incubation in MBL medium, and their concentration was quantified using a hemocytometer. (G) Germination rate. Conidial germination was observed by incubating 300 conidia for 2 or 8 hours on WA plates. Germination rates were compared, with different letters indicating significant differences based on ANOVA followed by the LSD test (*p*= 0.05).

### FgDML1 plays a critical role in both asexual and sexual reproduction

To examine the role of FgDML1 in asexual reproduction, a series of experiments were conducted on conidia. Morphological analysis revealed that conidia from the ΔFgDML1 mutant were thinner and longer than those from the PH-1 and ΔFgDML1*-C* (Fig. 2C). Furthermore, statistical analysis of septum number indicated a significant increase in conidia with ≥7 septa in the ΔFgDML1 compared to the controls (Fig. 2E). Conidial production was significantly reduced in ΔFgDML1, with fewer spores produced compared to PH-1 and ΔFgDML1-C (Fig. 2F). The germination rate of ΔFgDML1 conidia was also significantly lower than that of the controls (Fig. 2G). Regarding sexual reproduction, ΔFgDML1 failed to produce perithecia or ascospores on both the 7th and 14th days of incubation (Fig. 2D).

### FgDML1 is essential for virulence

To investigate the role of FgDML1 in virulence, pathogenicity assays were performed on wheat coleoptiles and wheat leaves. The results revealed that ΔFgDML1 caused only restricted infection at the inoculation site on wheat coleoptiles and failed to induce systemic infection, in contrast to PH-1 and ΔFgDML1-C, which exhibited extensive colonization (Fig. 3A, B). Similarly, ΔFgDML1 displayed a near-complete loss of virulence on wheat leaves (Fig. 3C, D). These findings demonstrate that FgDML1 is a critical positive regulator of virulence in *F. graminearum*.

**FIG. 3.**
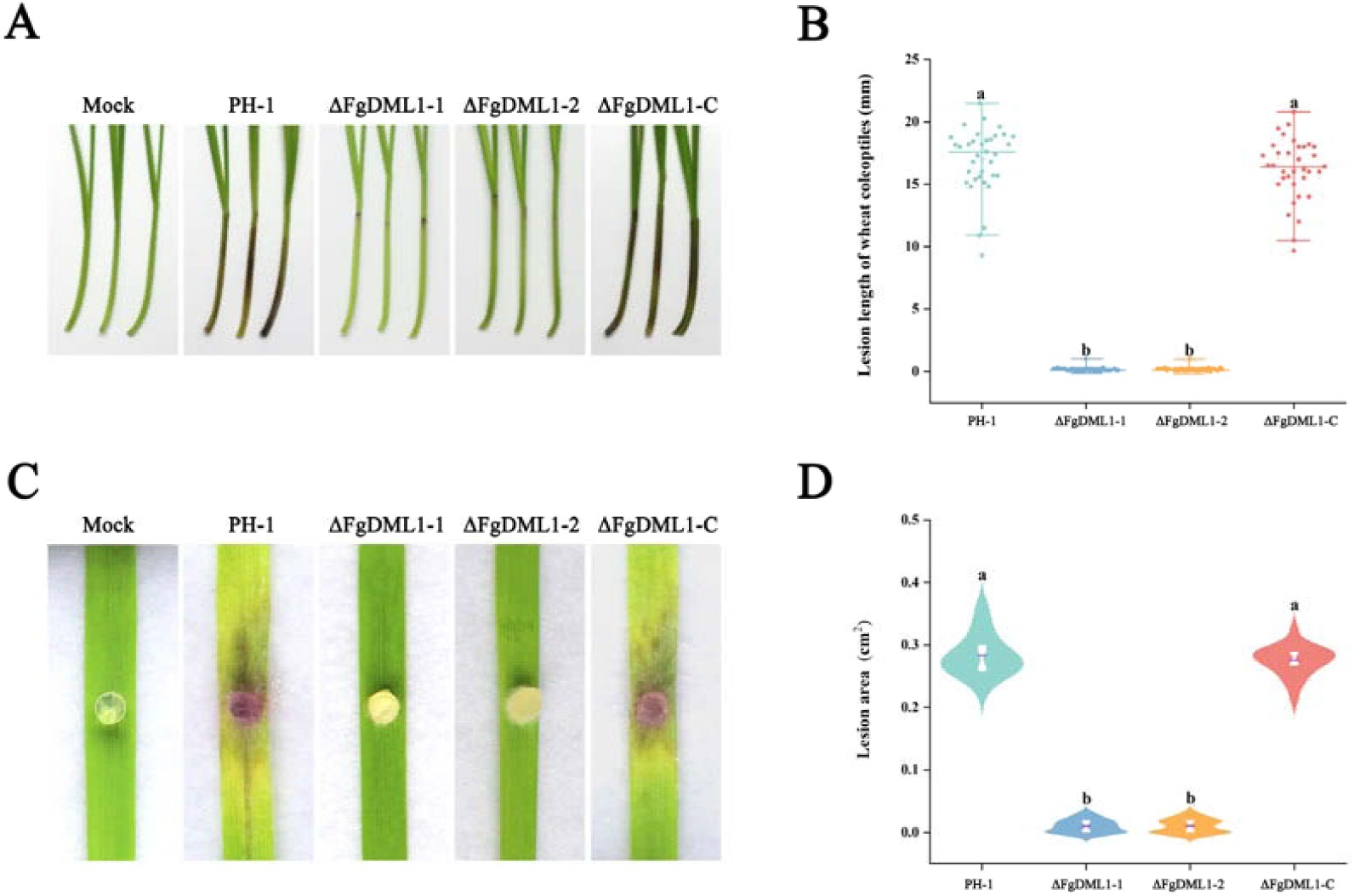
FgDML1 is required for both the virulence of *Fusarium graminearum*. (A) Lesion lengths on wheat coleoptiles caused by different strains. These images were taken at 14 dpi. (B) Dot plot showing the lesion lengths on wheat coleoptiles at 14 dpi tested with different strains. (C) Disease symptoms on wheat leaves inoculated with PH-1, ΔFgDML, and ΔFgDML-C. Images were taken at 5 dpi. (D) Disease area on wheat leaves inoculated with different strains at 5 dpi. Bars with the same letter indicate no significant difference according to a significant difference (LSD) test at *p* = 0.05.

### FgDML1 is essential for the biosynthesis of the DON toxin

To evaluate whether the deletion of FgDML1 influences DON production, DON levels were measured under toxin-inducing conditions (28°C, 145 *× g*, 7 days incubation). The results revealed that DON production in ΔFgDML1 was reduced by nearly 80% compared to PH-1 and significantly lower than that of both PH-1 and ΔFgDML1-C (Fig. 4A). To further elucidate the regulatory relationship between FgDML1 and DON biosynthesis, the transcriptional levels of *FgTri5* and *FgTri6* were analyzed and found to be significantly downregulated in ΔFgDML1 (Fig. 4B). Moreover, examination of toxisomes in the mutant demonstrated that their formation was severely impaired, with almost no detectable toxisomes in ΔFgDML1(Fig. 4C). Western blot analysis of the DON toxin marker protein FgTri1 further conirmed its markedly reduced levels in ΔFgDML1 compared to PH-1 (Fig. 4D). These findings suggest that FgDML1 is essential for regulating DON production in *F. graminearum*.

**FIG. 4.**
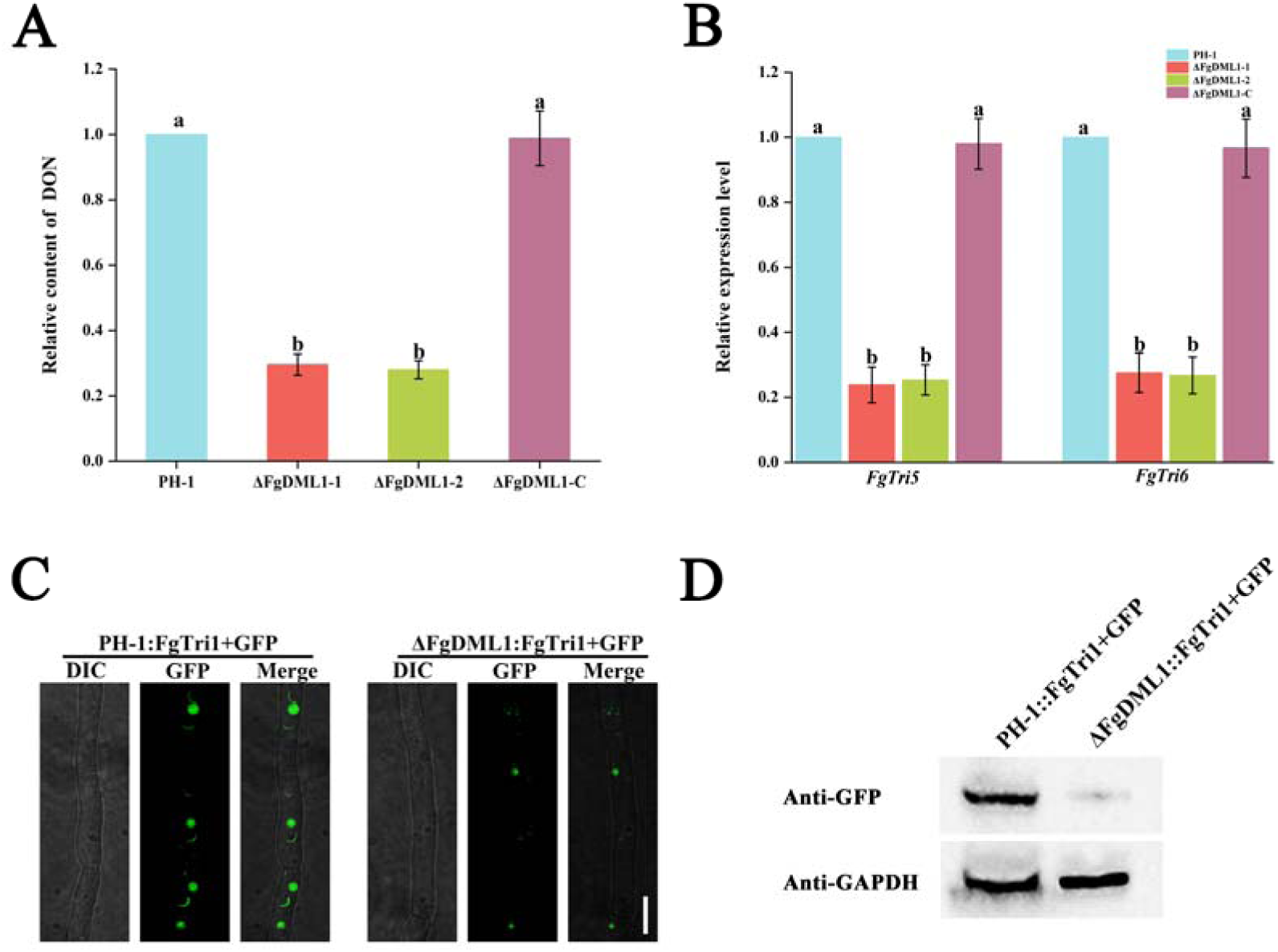
FgDML1 is required for the biosynthesis of DON toxin in *Fusarium graminearum*. (A) ΔFgDML1 exhibited decreased DON content. All strains were incubated in trichothecene biosynthesis induction (TBI) medium for 7 days. (B) Relative mRNA expression levels of *FgTRI5*, and *FgTRI6* in the tested strains. Mycelia were harvested after 2 days of cultivation in TBI medium, and mRNA were extracted for quantification using the 2^-ΔΔCT^ method. *FgGapdh* was used as the reference gene. (C) FgDML1 impaired the formation of DON toxisomes, as visualized by Tri1-GFP labeling of toxisomes. Strains were cultured in TBI for 36 hours. Bar = 10 μm (D) ΔFgDML1 exhibited reduced expression levels of Tri1-GFP. Strains were cultured in TBI for 36 hours, and mycelia were harvested for western blot analysis. Bars with the same letter indicate no significant difference according to a significant difference (LSD) test at *p* = 0.05.

### The deletion of FgDML1 results in reduced synthesis of both ATP and acetyl-CoA

The results showed that ATP levels in ΔFgDML1 were significantly lower compared to PH-1 and ΔFgDML1-C (Fig. 5D). Previous studies have established a positive correlation between ATP levels and acetyl-CoA. Consistent with this, acetyl-CoA levels in ΔFgDML1 were also significantly reduced compared to the controls (Fig. 5C). In conclusion, the deletion of FgDML1 leads to a reduction in both the key energy molecule ATP and the DON precursor acetyl-CoA.

**FIG. 5.**
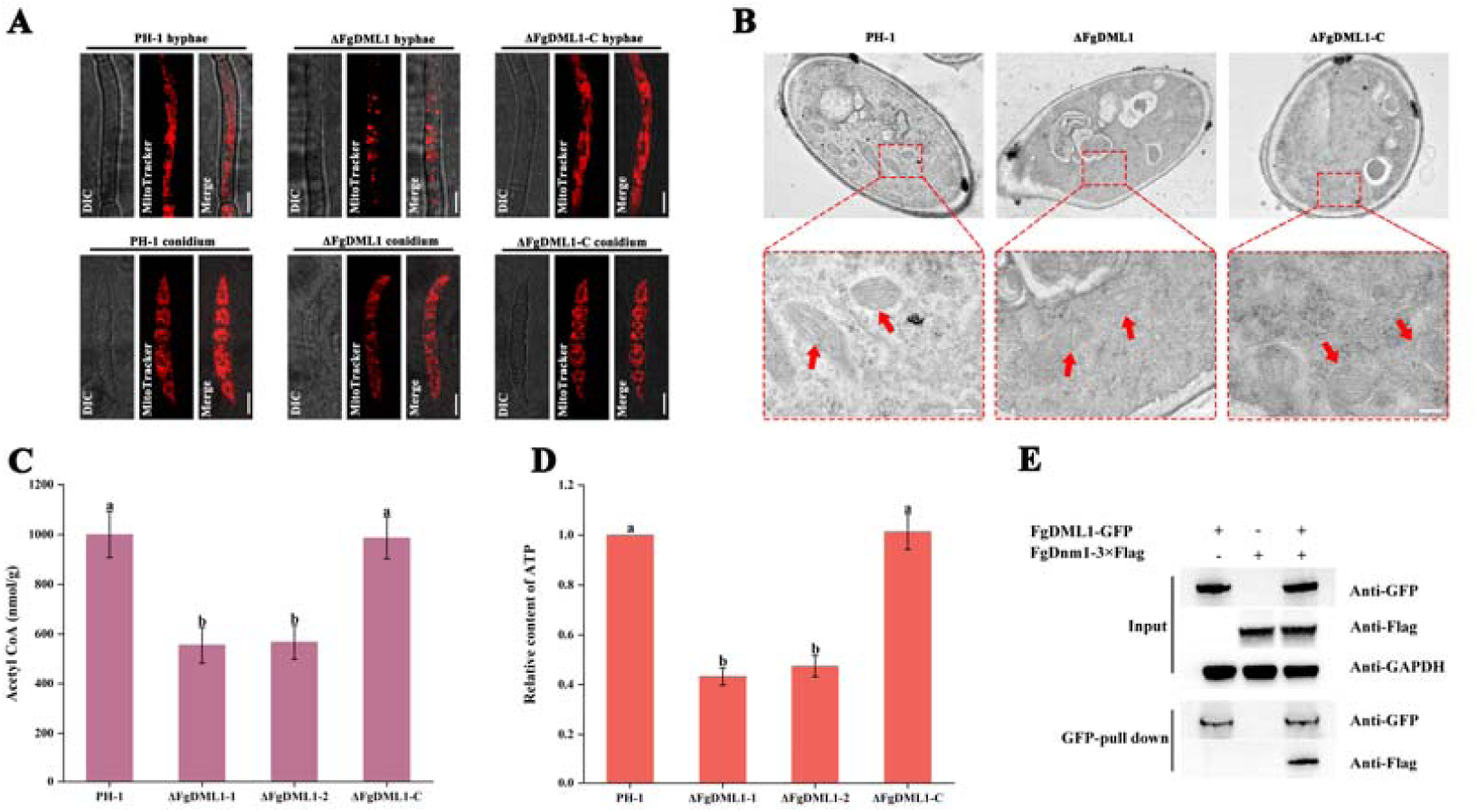
FgDML1 modulates mitochondrial morphology via interaction with FgDnm1 and negatively regulates Acetyl-CoA and ATP synthesis. (A) Effects of FgDML1 on mitochondrial morphology. Mycelial plugs of the wild-type strain PH-1, the ΔFgDML1 deletion mutant, and the complemented mutant ΔFgDML1-C were cultured in YEPD medium for 36 hours. Mitochondria were stained using a mitochondria-specific fluorescent dye and observed under a confocal microscope. Additionally, conidial germlings were cultured in MBL medium and subjected to the same staining and imaging procedures. Scale bar = 5 μm. (B) Ultrastructural analysis of mitochondria by transmission electron microscopy (TEM). Mitochondrial ultrastructure was further examined in PH-1, ΔFgDML1, and ΔFgDML1-C using TEM. Mitochondria are indicated by red arrows. Scale bar = 200 nm. (C) FgDML1 Deletion Reduces Acetyl-CoA Levels. Mycelial cultures of PH-1, ΔFgDML1, and ΔFgDML1-C were incubated in YEPD medium for 36 hours. Acetyl-CoA levels were quantified using a commercial assay kit, following the instructions provided by the commercial assay kit. (D) FgDML1 deletion reduces ATP levels. The ATP content of PH-1, ΔFgDML1, and ΔFgDML1-C was measured following the same culture conditions as in (C), using a commercial ATP assay kit. (E) Co-immunoprecipitation (Co-IP) confirms interaction between FgDML1 and FgDnm1. Total protein extracts from strains expressing tagged versions of FgDML1 and FgDnm1 were separated by SDS-PAGE and subjected to immunoblot analysis. Co-IP was performed using ChromoTek GFP-Trap® magnetic agarose to pull down interacting proteins. Immunodetection was carried out using polyclonal anti-Flag and monoclonal anti-GFP antibodies. Monoclonal anti-GAPDH antibody was used as a reference for protein samples. Bars with the same letter indicate no significant difference according to a significant difference (LSD) test at *p* = 0.05.

### FgDML1 regulates mitochondrial morphology by interacting with FgDnm1

To assess whether the loss of FgDML1 induces mitochondrial alterations, mitochondria-specific dyes were utilized, revealing fragmented mitochondria in the ΔFgDML1 mutant, whereas PH-1 and ΔFgDML1-C displayed a typical network-like mitochondrial distribution (Fig. 5A). Further analysis via Transmission Electron Microscope (TEM) identified damaged mitochondrial structures in the ΔFgDML1 mutant, in contrast to the well-preserved mitochondrial morphology observed in the control strains (Fig. 5B). Given that FgDnm1 is known to be a critical protein for mitochondrial fission, the interaction between FgDML1 and FgDnm1 was verified through co-immunoprecipitation (CoIP) assays (Fig. 5E). These findings indicate that FgDML1 plays a pivotal role in maintaining mitochondrial morphology by interacting with FgDnm1.

### The deletion of FgDML1 reduces Complex III activity, induces compensatory upregulation of FgQCR2, FgQCR8, FgQCR9

Deletion of FgDML1 reduced *F. graminearum* sensitivity to the Qi-site fungicide cyazofamid, while having no effect on sensitivity to the Qo-site inhibitors pyraclostrobin and trifloxystrobin, nor to other respiratory inhibitors such as pydiflumetofen, fluxapyroxad, and fluopyram (Fig. 6A, B) (Fig. S3). Measurement of Complex III activity in ΔFgDML1 showed a 38.28% reduction in enzyme activity compared to PH-1 (Fig. 6C). To further investigate the reduced sensitivity to cyazofamid, transcription levels of the three subunits (*FgCytb, FgCytC1, FgISP*) and five assembly factors (*FgQCR2, FgQCR6, FgQCR7, FgQCR8, FgQCR9*) of Complex III were analyzed. The results revealed that, upon Cyazofamid treatment, the expression of *FgQCR2*, *FgQCR7*, and *FgQCR8* was significantly upregulated in ΔFgDML1 (Fig. 6D). To further clarify whether the upregulated expression of *FgQCR2*, *FgQCR7*, and *FgQCR8* genes affects their protein expression levels, we measured the protein levels. The results showed that the protein expression levels of FgQCR2, FgQCR7, and FgQCR8 in ΔFgDML1 were higher than those in PH-1(Fig. 6F). Subsequently, we overexpressed *FgQCR2*, *FgQCR7*, and *FgQCR8* in the wild-type background, and the corresponding overexpression mutants exhibited reduced sensitivity to cyazofamid(Fig. 6E).

**FIG. 6.**
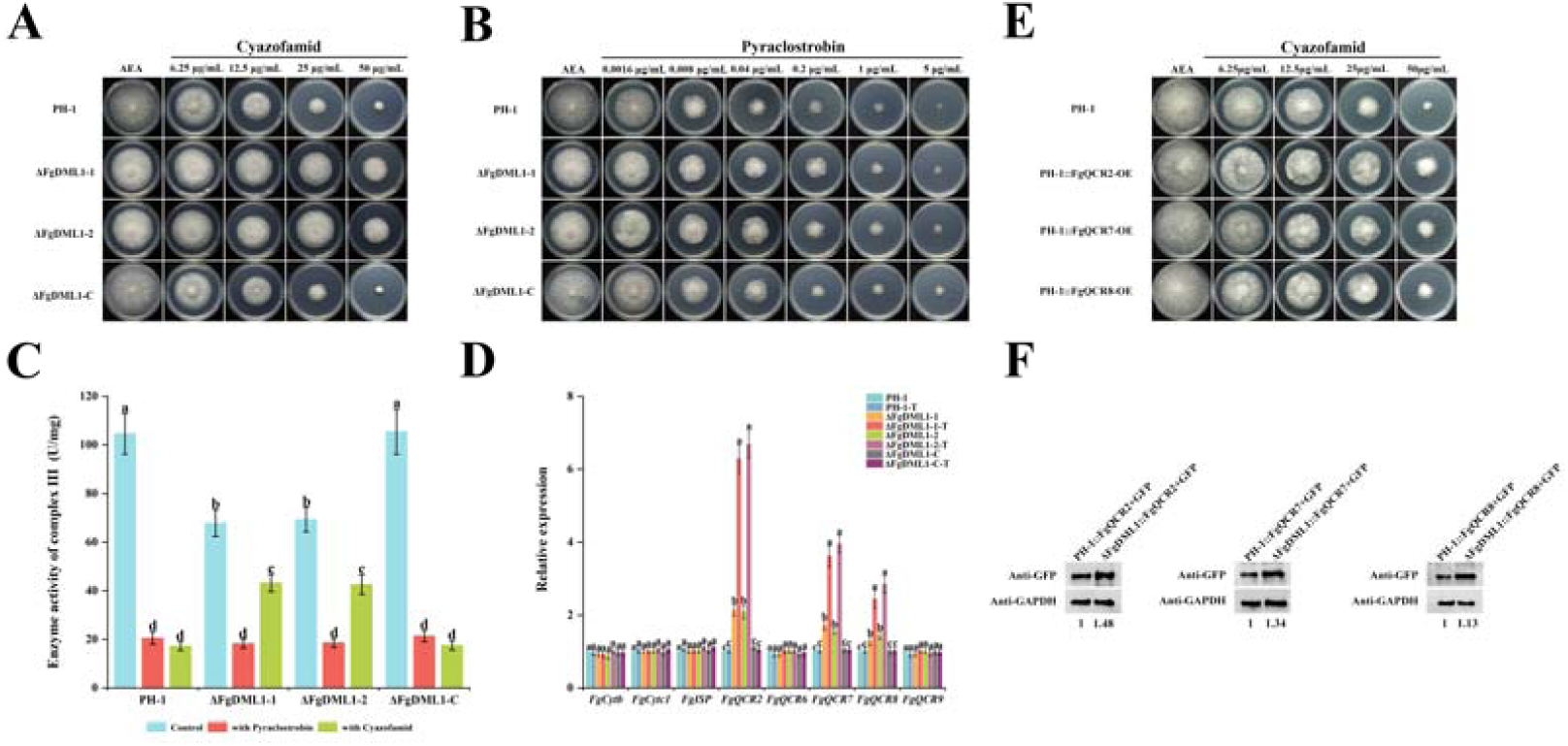
FgDML1 modulates the sensitivity of *Fusarium graminearum* to cyazofamid. (A) ΔFgDML1 exhibits reduced sensitivity to cyazofamid. PH-1, ΔFgDML1, and ΔFgDML1-C were cultured on AEA medium modified with cyazofamid, with 50 μg/mL SHAM included to inhibit the alternative oxidase pathway. (B) Deletion of *FgDML1* did not affect *F. graminearum* sensitivity to pyraclostrobin. The tested strains were incubated on AEA medium containing pyraclostrobin and 50 μg/mL SHAM to assess their response to the fungicide. (C) FgDML1 deletion reduces complex III enzymatic activity. Mycelial cultures of PH-1, ΔFgDML1, and ΔFgDML1-C were grown in YEPD for 36 hours, then treated with 30 μg/mL cyazofamid, 1.5 μg/mL pyraclostrobin, and 50 μg/mL SHAM for an additional 12 hours. The enzymatic activity of complex III was quantified using a commercial assay kit. (D) mRNA expression levels of complex III subunits and assembly factors in PH-1, ΔFgDML1, and ΔFgDML1-C. PH-1, ΔFgDML1, and ΔFgDML1-C were incubated in liquid AEA medium for 36 hours, followed by treatment with 30 μg/mL cyazofamid and 50 μg/mL SHAM. The relative mRNA expression levels of complex III subunits and assembly factors were quantified using the 2^-ΔΔCT^ method. "T" denotes cyazofamid treatment. Bars with the same letter indicate no significant difference according to a significant difference (LSD) test at *p* = 0.05. (E) Overexpression of *FgQCR2*, *FgQCR7*, and *FgQCR8*, respectively, in the PH-1 background resulted in reduced sensitivity to cyazofamid in the overexpression mutants. (F) Protein expression levels of FgQCR2, FgQCR7, and FgQCR8 in PH-1 and ΔFgDML1. Strains were cultured in AEA Liquid Medium for 36 h, and mycelia were harvested for western blot analysis.

### Overexpression of *FgQCR2*, *FgQCR8*, and *FgQCR9* may alters the conformation of the QI site, resulting in reduced sensitivity to cyazofamid

Since Complex III is involved in the action of both cyazofamid (targeting the QI site) and pyraclostrobin (targeting the QO site), the sensitivity of ΔFgDML1 to cyazofamid and pyraclostrobin was investigated. The results showed that the deletion of FgDML1 did not alter sensitivity to a pyraclostrobin (Fig. 6B). ΔFgDML1 was subsequently treated with pyraclostrobin or cyazofamid, and Complex III enzyme activity was measured. After pyraclostrobin treatment, no significant change in enzyme activity was observed in ΔFgDML1 compared to PH-1 and ΔFgDML1-C (Fig. 6C). However, when treated with cyazofamid, a significant increase in Complex III enzyme activity was observed in ΔFgDML1, while the control groups maintained relatively low enzyme activity (Fig. 6C). These findings, combined with the upregulation of assembly factors, suggest that the elevated expression of assembly factors alters the protein conformation at the QI site, modulating sensitivity to cyazofamid.

### FgDML1 regulates sensitivity to various stresses

*F. graminearum* encounters various stresses during infection plant, which significantly influence its ability to infect. To investigate its response to different stress conditions, the sensitivity of ΔFgDML1 to various environmental factors was assessed, including osmotic stress (NaCl, KCl, sorbitol), cell wall stress (Congo red), membrane stress (SDS), as well as high temperature (30°C) and low temperature (15°C) stresses. The results revealed that ΔFgDML1 displayed significantly reduced sensitivity to osmotic stress compared to PH-1 and ΔFgDML1-C (Fig. 7A, B). No significant difference was observed in sensitivity to membrane stress, while a notable change in sensitivity to cell wall stress was observed (Fig. 7C, D). Additionally, differences in response to both high and low temperature stress were noted in ΔFgDML1 when compared to the control strains PH-1 and ΔFgDML1-C (Fig. 7E, F).

**FIG. 7.**
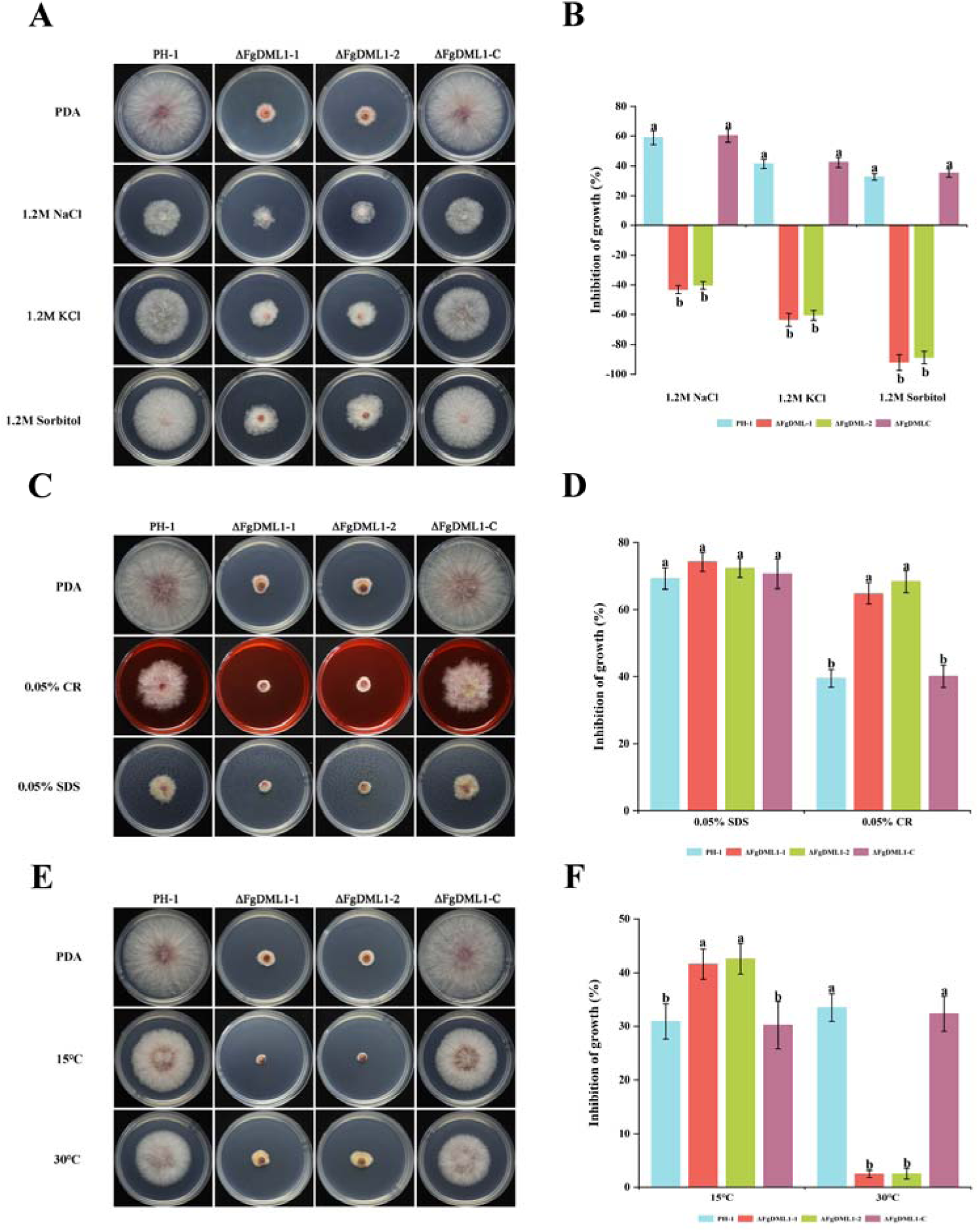
Response of the strains to different stresses. (A) Reduced sensitivity to osmotic stress in ΔFgDML1. PH-1, ΔFgDML1 and ΔFgDML1-C were subjected to osmotic stress using NaCl, KCl, and Sorbitol to evaluate their sensitivity. (B) Statistical analysis of growth inhibition under osmotic stress. Growth inhibition rates of all strains under NaCl, KCl, and Sorbitol stress were statistically analyzed. (C) Sensitivity to cell wall and membrane damaging agents. The tested strains were exposed to cell wall-damaging agent Congo Red (CR) and cell membrane-damaging agent Sodium dodecyl sulfate (SDS) to assess their sensitivity. (D) Statistical analysis of growth inhibition under CR and SDS stress. Growth inhibition rates of all strains under CR and SDS stress were statistically analyzed. (E) Sensitivity to temperature stresses. The sensitivity of the tested strains to different temperature stresses was evaluated. (F) Statistical analysis of growth inhibition under temperature stress. Growth inhibition rates of all strains under various temperature stresses were statistically analyzed. Significant differences were determined using the LSD test at *p* = 0.05. Bars with the same letter indicate no significant difference.

## Discussion

Misato Protein was originally identified in *Drosophila melanogaster*, where it was reported to play essential roles in mitochondrial DNA inheritance, cell division, and energy metabolism(Miklos et al., 1997). The Misato/DML1 protein family is evolutionarily conserved from yeast to humans and plays a critical role in mitochondrial regulation. In *S. cerevisiae*, *DML1* is an essential gene; its deletion is lethal, while its overexpression results in fragmented mitochondrial networks and aberrant cellular morphology, underscoring its necessity for normal mitochondrial function (Gurvitz et al., 2002). Similarly, in *Homo sapiens*, the homolog Misato localizes to the mitochondrial outer membrane, and both its depletion and overexpression are sufficient to disrupt mitochondrial morphology and distribution (Kimura and Okano, 2007). However, its functional characterization in filamentous fungi remains largely unexplored. Our study defines a novel mechanism through which FgDML1 governs mitochondrial homeostasis. We demonstrate that FgDML1 directly interacts with the key mitochondrial fission regulator FgDnm1 and positively modulates cellular bioenergetic metabolism, as evidenced by elevated ATP and acetyl-CoA levels (Fig. 8). In this study, we identified and analyzed the homolog FgDML1 in the phytopathogenic fungus *F. graminearum*. The FgDML1 protein contains conserved Misato_Tub_SegII and Tubulin_3 domains, with high homology observed among species within the *Fusarium* genus but significant genetic divergence from other taxa (Fig. 1A). In *S. cerevisiae*, DML1 is a critical regulator of mitochondrial fission and fusion dynamics, maintaining intracellular energy homeostasis by modulating mitochondrial morphology and distribution(Gurvitz et al., 2002). Our study demonstrated that FgDML1 localized to mitochondria in *F. graminearum* (Fig. 1B). Deletion mutants of *FgDML1* exhibited phenotypes indicative of mitochondrial dysfunction, including fragmented mitochondrial morphology (Fig. 5A). TEM further confirmed structural and dynamic damage to mitochondria in the ΔFgDML1 (Fig. 5B). Structural analysis suggests that DML1 shares similarities with GTPase family proteins, indicating potential GTPase activity(Gurvitz et al., 2002). GTPases primarily hydrolyze GTP, but their activity is intricately linked to ATP production and utilization. These enzymes regulate critical cellular processes such as energy metabolism, signal transduction, protein synthesis, and cytoskeletal remodeling, all of which influence ATP levels(Mrnjavac and Martin, 2025; Pb et al., 2023; Rogne et al., 2018). In agreement with this, we observed a significant reduction in ATP content in the FgDML1 deletion mutant, underscoring its role in mitochondrial energy metabolism (Fig. 5D). In *H. sapiens*, DML1 also localizes to mitochondria, where its silencing results in fragmented and aggregated mitochondria with reduced network continuity and fusion activity (Kimura and Okano, 2007). Our findings indicate that FgDML1 plays a conserved role in maintaining mitochondrial integrity and energy homeostasis. Notably, FgDML1 was found to interact with FgDnm1 (Fig. 5E), FgDnm1 is a key dynamin-related protein mediating mitochondrial fission(Griffin et al., 2005; Kang et al., 2023), suggesting that FgDML1 may form a complex with FgDnm1 to regulate mitochondrial fission and fusion processes. To our knowledge, this is the first report documenting an interaction between DML1 and Dnm in any fungal species, including model organisms such as *S. cerevisiae*. This novel finding provides new insights into the molecular mechanisms underlying mitochondrial dynamics in filamentous fungi.

**FIG. 8.**
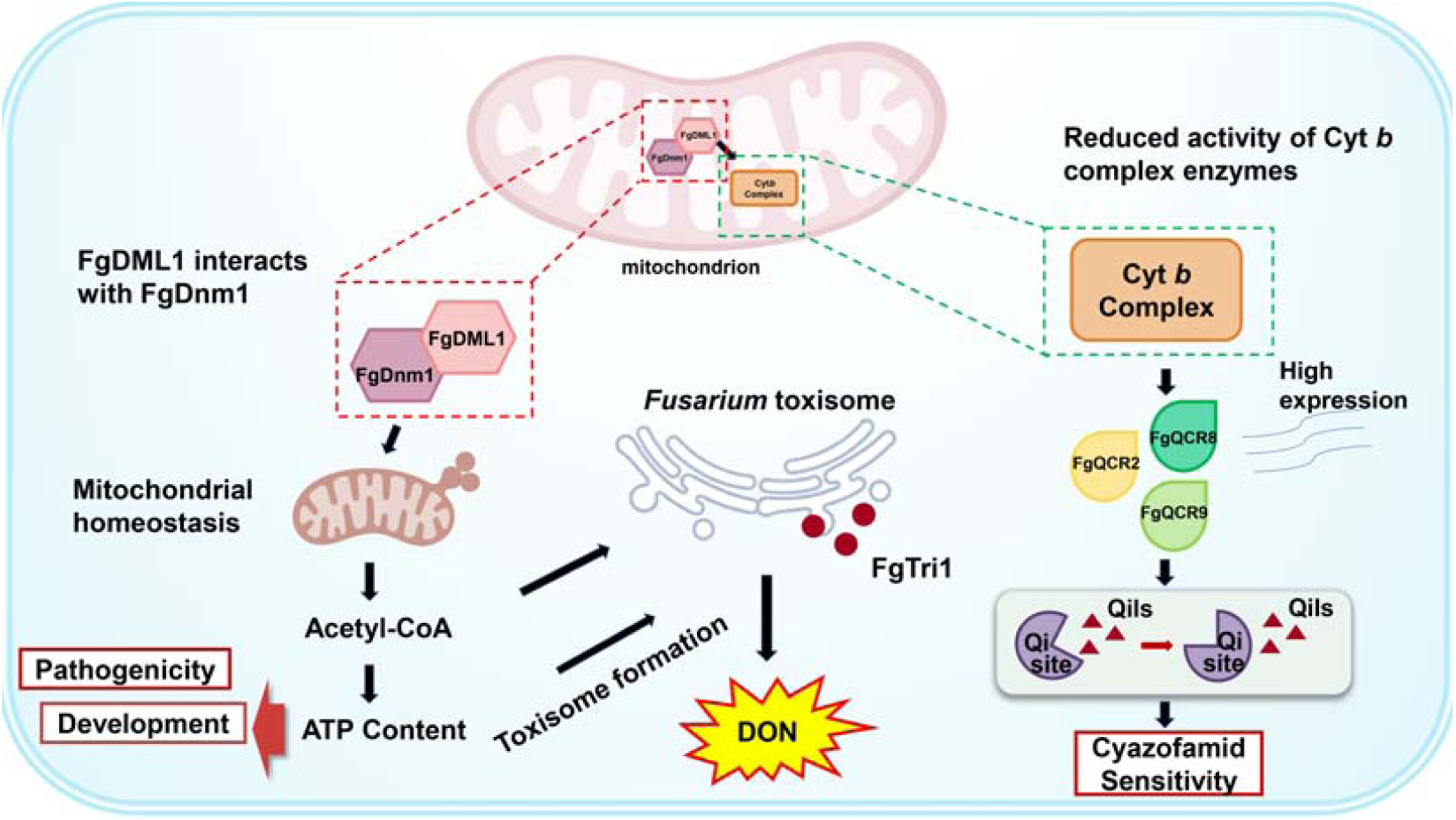
Model of FgDML1 regulation of *Fusarium graminearum* sensitivity to DON toxin synthesis and cyazofamid fungicide. FgDML1 interacts with FgDnm1 to modulate mitochondrial morphology, thereby positively regulating acetyl-CoA and ATP levels, which influence toxisome formation and ultimately affect DON production. Loss of FgDML1 reduces complex III enzymatic activity, leading to the upregulation of the assembly factors *FgQCR2*, *FgQCR8*, and *FgQCR9*. This alteration in assembly factor expression changes the conformation of the Qi site, leading to reduced sensitivity to cyazofamid.

Secondary metabolite biosynthesis is generally regarded as an energy-intensive process that is tightly coupled to cellular energy metabolism. ATP serves as the primary energy currency supporting enzymatic reactions, macromolecule synthesis, and subcellular organization required for secondary metabolism. Disruption of ATP generation has been shown to directly impair toxin biosynthesis: for example, silencing of ATP synthase subunit α (AtpA) significantly reduces ATP synthesis and inhibits the production of the TcdA and TcdB toxins(Marreddy et al., 2024). Similarly, in plants, ATP depletion leads to a metabolic shift in which growth and basic physiological processes are prioritized at the expense of energetically costly secondary metabolites, including toxins(Xiao et al., 2024). Together, these findings highlight ATP availability as a key determinant of secondary metabolite production across biological systems.

In filamentous fungi, mitochondria play a central role in sustaining cellular ATP levels through oxidative phosphorylation and are therefore critical for biosynthetic and stress-adaptive processes. In *F. graminearum*, mutants defective in mitochondrial components, such as the voltage-dependent anion channel (mitochondrial porin), exhibit aberrant mitochondrial morphology, reduced ATP production, and markedly decreased DON accumulation and virulence (Han et al., 2022). These observations establish a direct link between mitochondrial energy metabolism and secondary metabolite output, supporting the notion that intact mitochondrial function and adequate ATP supply are prerequisites for robust DON production.

Consistent with this energy-dependent framework, biosynthesis of the mycotoxin DON in *F. graminearum* requires substantial ATP input. In the present study, ATP content in the ΔFgDML1 mutant was significantly lower than in the wild-type PH-1 and the complemented strain ΔFgDML1-C, and DON production was concomitantly reduced (Fig. 4A). Importantly, DON levels were normalized to mycelial dry weight, indicating that the observed reduction reflects a decreased biosynthetic capacity per unit biomass rather than a secondary consequence of reduced fungal growth. This distinction demonstrates that impaired DON production in the ΔFgDML1 mutant arises primarily from metabolic limitations.

At the cellular level, ATP depletion compromises multiple energy-dependent steps required for DON biosynthesis. The formation of toxisomes, which are specialized subcellular structures responsible for the spatial organization of DON biosynthetic enzymes, is essential for efficient mycotoxin production and is an ATP-dependent process. Reduced ATP levels disrupt toxisome assembly, and accordingly, the ΔFgDML1 mutant was unable to form functional toxisomes (Fig. 4C). In parallel, western blot analysis revealed a marked reduction in the abundance of the DON biosynthetic enzyme FgTri1 (Fig. 4D). In addition, ATP-dependent processes are directly involved in the biogenesis of the DON biosynthetic machinery: the ATPase activity of myosin I (FgMyo1) is required for efficient translation of key DON biosynthetic enzymes, and disruption of its ATPase function results in reduced DON production(Tang et al., 2018). These findings further underscore the dependence of DON biosynthesis on cellular energy status.

DON production is also regulated at the transcriptional level by the TRI gene cluster, with Tri5 and Tri6 serving as core components of the biosynthetic pathway. Tri5 encodes trichodiene synthase, which catalyzes the first committed step of DON biosynthesis. In the ΔFgDML1 mutant, expression levels of FgTri5 and FgTri6 were significantly downregulated (Fig. 4B), suggesting that impaired energy metabolism indirectly affects transcription of DON biosynthetic genes. Although no direct regulatory role of DML family proteins in gene expression has been reported in Saccharomyces cerevisiae or Drosophila melanogaster, their established functions in cell division and microtubule organization raise the possibility that FgDML1 indirectly influences gene expression through effects on chromatin organization or cell-cycle progression(Schulze and Wallrath, 2007).

In addition to reduced ATP levels, deletion of FgDML1 resulted in a significant decrease in acetyl-CoA content (Fig. 5C), a key precursor for trichothecene biosynthesis. Acetyl-CoA links central carbon metabolism with secondary metabolite production, and its depletion further constrains DON biosynthesis by limiting substrate availability. Broader metabolomic studies support this relationship, showing that perturbations in TCA cycle intermediates and central carbon metabolism are closely associated with altered DON production, reinforcing a mechanistic linkage between energy generation and toxin biosynthesis(Atanasova-Penichon et al., 2018).

Taken together, these results support a model in which FgDML1 influences DON production indirectly by maintaining mitochondrial energy metabolism. Reduced ATP availability in the ΔFgDML1 mutant restricts energy-dependent biosynthetic processes, disrupts toxisome formation, diminishes DON biosynthetic enzyme abundance and gene expression, and limits precursor supply, ultimately leading to a substantial reduction in DON biosynthesis that is independent of fungal biomass effects.

*F. graminearum* is a highly destructive pathogen that poses a significant threat to wheat production worldwide. Cyazofamid is an effective fungicide commonly used to control plant pathogenic fungi. Cyazofamid specifically targets the Qi site of complex III in the mitochondrial respiratory chain(Bolgunas et al., 2006; Mitani et al., 2001). Complex III, also known as the cytochrome bc1 complex, plays a critical role in the electron transport chain by transferring electrons from ubiquinone to cytochrome c and pumping protons from the mitochondrial matrix into the intermembrane space(Fernández-Ortuño et al., 2008; Sarewicz et al., 2013). This complex consists of several core subunits, including cytochrome b, cytochrome c1, and iron-sulfur proteins, as well as multiple assembly factors. It contains two quinone binding sites: the Qo site (quinone oxidation site) and the Qi site (quinone reduction site)(Vercellino and Sazanov, 2022; Zara et al., 2009). Numerous fungicides have been developed to inhibit the Qo site (e.g., pyraclostrobin, azoxystrobin)(Nuwamanya et al., 2022; Peng et al., 2022) and the Qi site (e.g., cyazofamid)(Mitani et al., 2001) of the cytochrome bc1 complex. In our study, we observed that the deletion of *FgDML1* significantly reduced the sensitivity of *F. graminearum* to cyazofamid (Fig. 6A), while its sensitivity to pyraclostrobin remained unaffected (Fig. 6B). In the ΔFgDML1 mutant, we detected a decrease in complex III enzyme activity (Fig. 6C) and noted the upregulation of assembly factors *FgQCR2*, *FgQCR7*, and *FgQCR8* following cyazofamid treatment (Fig. 6D). Further enzyme activity assays revealed that after pyraclostrobin treatment, ΔFgDML1 exhibited a 73.06% reduction in complex III activity, compared to an 80.38% decrease in the wild-type strain PH-1 (Fig. 6C). Interestingly, after cyazofamid treatment, the decrease in enzyme activity in ΔFgDML1 was only 36.31%, while PH-1 showed a dramatic 83.46% reduction in complex III activity. *FgQCR2*, *FgQCR7*, and *FgQCR8* are critical assembly factors that play essential roles in the proper assembly of complex III(Zhan et al., 2024). These factors stabilize the core subunit structure, promoting correct folding and functional operation of complex III, particularly during the assembly of the Qi site. This ensures the proper formation and function of the Qi site, thus supporting the normal operation of the mitochondrial respiratory chain(Wang et al., 2024; Zara et al., 2007). Previous studies have reported that mutations in the Complex III assembly factors TTC19, UQCC2, and UQCC3 impair the assembly and activity of Complex III (Feichtinger et al., 2017; Wanschers et al., 2014). Given the role of FgDML1 in mitochondrial homeostasis, we further evaluated the sensitivity of the ΔFgDML1 mutant to other commonly used respiratory chain inhibitors, including pydiflumetofen, fluxapyroxad, fluopyram, and trifloxystrobin. The results indicated that FgDML1 deletion had no impact on the sensitivity of *F. graminearum* to these fungicides (Fig. S3). In conclusion, our findings suggest that the overexpression of assembly factors *FgQCR2*, *FgQCR7*, and *FgQCR8* in ΔFgDML1 potentially modifies the conformation of the Qi site, which specifically modulates the sensitivity of *F. graminearum* to cyazofamid.

In *H. sapiens*, mutations or deletions in DML1 are associated with mitochondrial diseases characterized by early-onset myopathy and cerebellar ataxia(Chen et al., 2022; Nasca et al., 2017). In our study, the deletion of *FgDML1* in *F. graminearum* resulted in significant defects in vegetative growth, including a markedly reduced growth rate (Fig. 2A) and an aberrant, excessively curved hyphal morphology (Fig. 2B). Notably, the loss of FgDML1 rendered *F. graminearum* incapable of completing sexual reproduction (Fig. 2D), thereby potentially reducing its primary inoculum sources. Additionally, the deletion of FgDML1 impaired asexual reproduction and pathogenicity (Fig. 2) (Fig. 3), critical factors for fungal survival and host infection. Furthermore, the ΔFgDML1 mutant exhibited altered responses to various stress conditions (Fig. 7), suggesting a broader role for FgDML1 in regulating fungal stress tolerance.

In the present study, FgDML1 is localized to the mitochondria and interacted with FgDnm1, a key regulator of mitochondrial morphology, positively regulates ATP and acetyl-CoA levels, influences toxin body formation, and ultimately leads to a significant reduction in DON toxin biosynthesis. Furthermore, deletion of FgDML1 impairs the enzymatic activity of Complex III, which subsequently induces the upregulation of the assembly factors FgQCR2, FgQCR8, and FgQCR9, resulting in conformational changes at the Qi site of Complex III and decreased sensitivity to cyazofamid. In addition, the absence of FgDML1 has profound effects on pathogenicity, fungal development, and mitochondrial morphology. Based on these findings, we propose a working model delineating the role of FgDML1 in regulating DON toxin biosynthesis and cyazofamid sensitivity in *F. graminearum* (Fig. 8). This study not only provides a theoretical basis for understanding the regulatory mechanisms underlying toxin biosynthesis in *F. graminearum* but also identifies potential molecular targets for the development of novel agents to control FHB.

## Material and methods

### Strains, culture conditions, and fungicides

The wild-type *F. graminearum* strain PH-1 was used as the parental strain for protoplast transformation experiments. All strains were preserved at -80°C. The tested strains were cultured in the dark on potato dextrose agar (PDA) at 25°C. To evaluate vegetative growth, complete medium (CM), minimal medium (MM), and V8 Juice Agar (V8) media were prepared as described previously(Tang et al., 2020). Mung bean liquid (MBL) medium was used for conidial production, while carrot agar (CA) medium was utilized to assess sexual reproduction(Wang et al., 2011). DON toxin production was measured using trichothecene biosynthesis-inducing (TBI) medium(Wang et al., 2024).

Cyazofamid (98%, BASF China Ltd.), pyraclostrobin (98%, BASF China Ltd.) pydiflumetofen (98%, China Syngenta crop Protection Co., Ltd.), trifloxystrobin (96%, Bayer Crop Science China Co., Ltd.), fluxapyroxad (98%, BASF China Ltd.) and fluopyram (96%, Bayer Crop Science China Co., Ltd.) were dissolved in dimethyl sulfoxide (DMSO) at stock concentrations of 50,000 μg/mL and 10,000 μg/mL, respectively.

### Generation of deletion mutants and complemented mutants

The split-marker method was employed to generate the ΔFgDML1 mutant(Yu et al., 2004). Specifically, approximately 1.2 kb of the upstream and downstream flanking regions of FgDML1 were amplified and fused with a dual-selection marker containing the hygromycin phosphotransferase (*HPH*) gene and the herpes simplex virus thymidine kinase (*HSV-tk*) gene fragments. The resulting fusion fragment was transformed into the wild-type *F. graminearum* PH-1 strain via polyethylene glycol (PEG)-mediated protoplast transformation. Transformants were selected on PDA plates containing either 100 μg/mL Hygromycin B (Yeasen, Shanghai, China) or 0.2 μmol/mL 5-Fluorouracil 2’-deoxyriboside (F2du) (Solarbio, Beijing, China)(Zhao et al., 2022). Positive transformants were initially screened by PCR and subsequently verified through Southern blot analysis. To generate the complemented mutant ΔFgDML1-C, the full-length coding sequence (CDS) of FgDML1 was introduced into the ΔFgDML1 mutant, replacing the dual-selection cassette. Furthermore, FgDML1-GFP and FgDnm1-3×Flag mutants were constructed using previously established methods for further functional studies(Wu et al., 2022). Specifically, FgDML1, including its native promoter region and open reading frame (ORF) (excluding the stop codon), was amplified.The PCR product was then fused with the *Xho*I-digested pYF11 vector. After transformation into E. coli and sequence verification, the plasmid was extracted and subsequently introduced into PH-1 protoplasts. For FgDnm1-3×Flag, the 3×Flag tag was added to the C-terminus of FgDnm1 by PCR, fused with the hygromycin resistance gene and the FgDnm1 downstream arm, and then introduced into PH-1 protoplasts. The overexpression mutant was constructed according to a previously described method. Specifically, the ORF of FgDML1 was amplified and the PCR product was ligated into the *Sac*II-digested pSXS overexpression vector. The resulting plasmid was then transformed into PH-1 protoplasts (Shi et al., 2023). For the construction of PH-1::FgTri1+GFP and ΔFgDML1::FgTri1+GFP, the ORF of FgTri1 was amplified and ligated into the *Xho*I-digested pYF11 vector as described above. The resulting vectors were then transformed into protoplasts of PH-1 or ΔFgDML1, respectively.

### Vegetative growth and conidiation assays

Fresh mycelial plugs of the tested strains were transferred to PDA, CM, MM, or V8 plates to evaluate mycelial growth rates. Conidial production was induced by incubating the strains in MBL medium, and the number of conidia was determined using a hemocytometer. The septa of conidia were visualized through calcofluor white (CFW) staining. For germination rate assessment, conidia were collected and inoculated onto water agar (WA) plates, followed by incubation at 25°C for 2 and 8 hours. Germination rates were then recorded at the respective time points. For each sample, a total of 200 conidia were counted. The experiment included three biological replicates with three technical replicates each.

### Sexual reproduction assay

Sexual reproduction experiments were performed on CA medium. Test strains were incubated on CA medium in complete darkness for 6 days. After incubation, the surface mycelia were removed using a mycelial scraper and 2.5% Tween 60 solution. The plates were then further incubated at 25°C under a 12-hour light/12-hour dark cycle to induce sexual development. Perithecia and ascospores were observed and recorded on days 7 and 14 using a Nikon SMZ25 fluorescent stereomicroscope and an Olympus IX-71 inverted fluorescent microscope. Each treatment group contains three biological replicates.

### Microscopic observation

To determine the subcellular localization of FgDML1, the FgDML1-GFP mutant was incubated in YEPD medium at 25°C with shaking at 145 × g for 36 hours. Subcellular localization was analyzed using a Leica TCS SP8 confocal microscope, with MitoTracker Red CMXRos (Yeasen, Shanghai, China) applied to confirm mitochondrial targeting through colocalization of fluorescence signals. To investigate toxisome morphology, the ΔFgDML1::Tri1+GFP mutant was cultured in TBI medium at 28°C with shaking for 36 hours, and the structural dynamics of toxisomes within the mycelium were visualized using Tri1-GFP fluorescence under confocal microscopy. Additionally, mitochondrial morphology was evaluated by culturing PH-1, ΔFgDML1, and ΔFgDML1-C in YEPD medium under identical conditions, followed by staining with MitoTracker Red CMXRos and confocal microscopy analysis to assess mitochondrial structural alterations, distribution, and dynamics. For ultrastructural observations, mycelial samples were fixed and dehydrated according to standard protocols, embedded, sectioned, and stained with uranyl acetate and lead citrate. Mitochondrial ultrastructure was examined using a Tecnai G2 Spirit Bio electron microscope (Thermo, MA, USA) at an accelerating voltage of 80 kV to compare the structural integrity and morphological differences among the strains. Each treatment group contains three biological replicates and three technical replicates.

### Determination of DON production and pathogenicity

As previously described (Tang et al., 2020; Wang et al., 2025), Specifically, coleoptiles were inoculated with conidial suspensions and incubated for 14 days, while leaves were inoculated with fresh mycelial plugs and incubated for 5 days, followed by observation and quantification of disease symptoms. DON toxin was measured using a Wise Science ELISA-based kit (Wise Science, Jiangsu, China) (Li et al., 2019; Zheng et al., 2018). Under toxin-producing conditions (28 °C, 145 rpm), fungal strains were cultured in TBI medium for 7 days. Cultures were initiated using freshly grown mycelia. After incubation, mycelia and culture filtrates were separated by filtration. The culture filtrates were collected for DON determination, while the mycelia were harvested for biomass analysis. The collected mycelia were washed with sterile distilled water and dried at 60 °C to constant weight. The dry weight of mycelia was recorded and used for normalization of DON production. One mycelial unit was defined as 1 g of dry mycelial biomass. DON concentration in the culture filtrates was quantified using an enzyme-linked immunosorbent assay (ELISA). Briefly, 50 μL of culture filtrate or DON standard solution was added to wells of a 96-well microplate pre-coated with DON antigen, followed by the addition of enzyme conjugate and antibody working solution according to the manufacturer’s instructions. After incubation and washing, color development was achieved using substrate solution and terminated by stop solution. Absorbance was measured at 450 nm using a microplate reader. A standard curve was generated using log -transformed DON concentrations of the standards and the corresponding percentage absorbance values. DON concentrations in the samples were calculated based on the standard curve. Total DON production was calculated according to the culture volume (30 mL) and subsequently normalized to mycelial dry weight. DON production was expressed as μg DON per g dry mycelium. Each treatment group contains three biological replicates and three technical replicates.

### Determination of acetyl-CoA content and ATP concentration

ATP content and acetyl-CoA levels were determined using the methods described previously(Hu et al., 2023; Song et al., 2024). The test strains were cultured in yeast extract peptone dextrose (YEPD) medium at 25°C and 145 × g for 36 hours. Mycelia were collected by filtration through three layers of lens-cleaning paper, and excess moisture was removed with filter paper. Subsequently, ATP and acetyl-CoA levels were quantified using ATP and acetyl-CoA assay kits (Solarbio, Beijing, China), respectively. Each treatment group contains three biological replicates and three technical replicates.

### Sensitivity determination

To investigate the sensitivity of ΔFgDML1 to QoI and QiI fungicides, the wild-type strain PH-1, ΔFgDML1 mutant, and the complemented mutant ΔFgDML1-C were inoculated on AEA plates supplemented with gradient concentrations of cyazofamid and pyraclostrobin. The wild-type PH-1 and its overexpression mutants (PH-1::FgQCR2-OE, PH-1::FgQCR7-OE, and PH-1::FgQCR8-OE) were assayed on AEA plates amended with cyazofamid and supplemented with SHAM. All strains were incubated at 25°C in darkness; however, due to ΔFgDML1 slower growth, the ΔFgDML1 mutant required a 5-day incubation period compared to the 3 days used for PH-1 and ΔFgDML1-C. The concentration gradients for each fungicide in the sensitivity assays were set up according to Supplementary Table S2. To inhibit the alternative respiratory pathway, 50 μg/mL SHAM was applied. The percentage inhibition (%) of mycelial growth was calculated using the formula: (mean control diameter - mean diameter at fungicide concentration) / (mean control diameter - 5 mm mycelial plug) × 100. To evaluate the sensitivity of ΔFgDML1 to various stress conditions, the test strains were cultured on PDA plates supplemented with stress-inducing agents, including NaCl, KCl, sorbitol, Congo red, and SDS. Plates were incubated at 25°C for 3 days, and colony growth was assessed. Each treatment group contains three biological replicates.

### Measurement of complex III enzyme activity

The enzyme activity of Complex III was determined in the PH-1, ΔFgDML1, and ΔFgDML1-C. Fresh mycelial plugs from each strain were inoculated into liquid AEA medium and incubated at 25°C with shaking at 145 × *g* for 36 hours. Subsequently, cyazofamid and pyraclostrobin were added to final concentrations of 30 μg/mL and 1.5 μg/mL, respectively, along with 50 μg/mL SHAM. After an additional 12-hour incubation, the mycelia were harvested, and Complex III activity was measured according to the complex III enzyme activity kit (Solarbio, Beijing, China) instructions. Briefly, 0.1 g of mycelia was homogenized with 1 mL of extraction buffer in an ice bath. The homogenate was centrifuged at 600 ×g for 10 min at 4°C. The resulting supernatant was then subjected to a second centrifugation at 11,100 ×g for 10 min at 4°C. The pellet was resuspended in 200 μL of extraction buffer and disrupted by ultrasonication (200 W, 5 s pulses with 10 s intervals, 15 cycles). Complex III enzyme activity was finally measured by adding the working solution as per the manufacturer’s protocol. Each treatment group contains three biological replicates and three technical replicates.

### RNA extraction and RT-qPCR assays

The test strains were transferred to liquid AEA medium and incubated at 25°C for 48 hours, with cyazofamid added to a final concentration of 20 µg/mL. The control group was not treated with any fungicide. To measure the transcriptional levels of genes involved in DON biosynthesis, the test strains were transferred to TBI medium and incubated for 3 days. Mycelium was collected and total RNA was extracted following the instructions provided by the Total RNA Extraction Kit (Tiangen, Beijing, China). cDNA synthesis was performed using Evo M-MLV Mix Kit (Accurate, Hunan, China). reverse transcription. Reverse Transcription Quantitative Polymerase Chain Reaction (RT-qPCR) was carried out using the QuantStudio 6 Flex real-time PCR system (Thermo, Fisher Scientific, USA) to assess the relative expression of three subunits of Complex III (*FgCytb*, *FgCytc1*, *FgISP*), five assembly factors (*FgQCR2*, *FgQCR6*, *FgQCR7*, *FgQCR8*, *FgQCR9*), and DON biosynthesis-related genes (*FgTri5* and *FgTri6*). The *FgCox1* gene was used as the reference gene for *FgCytb*, while *FgGapdh* was used as the reference for the other genes. Relative transcript levels were calculated using the 2^-ΔΔCT^ method. Each treatment group contains three biological replicates and three technical replicates.

### Western Blot assay and Co-IP assay

Total protein extraction was performed according to previously established protocols, and the extracted proteins were separated using sodium dodecyl sulfate-polyacrylamide gel electrophoresis (SDS-PAGE). Subsequently, proteins were transferred onto an Immobilon-P transfer membrane (Millipore, Billerica, USA). The membrane was probed with GFP antibody D191040 (BBI, Shanghai, China) and Flag antibody 30501ES60 (Yeasen, Shanghai, China). The FgDnm1-3×Flag fragment was introduced into PH-1 and FgDML1+GFP protoplasts, respectively, to obtain single-tagged and double-tagged strains. All tagged mutant strains were subjected to triple validation, including PCR, sequencing, and Western blot analysis. Total protein from tagged strains was incubated with ChromoTek GFP-Trap® Magnetic Agarose (ChromoTek, Planegg-Martinsried, Germany) beads. Proteins eluted from the beads were further co-incubated with GFP or Flag antibodies for detection. Gapdh antibody 60004-1-Ig (Proteintech, Rosemont, USA) was used as an internal control for protein normalization. Chemiluminescent detection was performed following incubation with the corresponding secondary antibody. All antibodies were diluted as follows: primary antibodies at 1:1000 and secondary antibodies at 1:10000. The experiment was independently repeated three times.

### Statistical analysis

All data were analyzed using one-way analysis of variance (ANOVA), followed by the least significant difference (LSD) test with a significance threshold of *p* < 0.05. The EC_50_ values were calculated using DPS statistical analysis software. Each experiment was independently repeated three times.

## Acknowledgements

This study was sponsored by National Natural Science Foundation of China (32272585) and National Key Research and Development Program of China (2022YFD1400900).

## CRediT authorship contribution statement

Chenguang Wang: methodology, investigation, editing, and writing original draft. Xuewei Mao: formal analysis, data curation. Weiwei Cong, Lin Yang.: performing the partial work of the experiments and analyzing the data. Yiping Hou: conceptualization, resources, writing−reviewing, supervision, funding acquisition.

